# Bioluminescent sentinel plants enable autonomous diagnostics of viral infections

**DOI:** 10.1101/2025.08.05.668616

**Authors:** Camilo Calvache, Marta Rodriguez-Rodriguez, Victor Vazquez-Vilriales, Elena Garcia-Perez, Aubin Fleiss, Mustafá Ezzeddin-Ayoub, Fabio Pasin, José Antonio Daròs, Karen S. Sarkisyan, Diego Orzaez, Marta Vazquez-Vilar

## Abstract

Plants engineered with synthetic genetic programs can transform how we monitor and manage the extension of crop pests and diseases. Here, we establish a bioluminescent platform in *Nicotiana benthamiana* for autonomous viral sensing based on the fungal bioluminescence pathway (FBP). We first demonstrate that recombinant viruses can deliver missing pathway components, enabling spatially resolved tracking of infection dynamics. Leveraging this starting point, we developed a dual-output sentinel circuit that uses a protease-responsive Bioluminescence Resonance Energy Transfer (BRET) module to report infection through a virus-triggered spectral shift in luminescence. In the absence of infection, plants emit a stable yellow glow indicating system integrity. Upon infection with potyviruses, cleavage of the BRET fusion by the virus-encoded NIa-Pro protease activates a distinct colour change detectable with low-cost imaging. This modular design is compatible with other pathogens carrying specific proteases and supports future multiplexing strategies. Our results highlight the potential of synthetic sentinel gene circuits as autonomous biosensors for precision crop protection.

## Introduction

The integration of continuous sensing technologies into production systems enables real-time monitoring of plant health, facilitating data-driven decision making and optimized crop management^1^. Synthetic biology offers a powerful framework for equipping plants with novel functions, including the ability to sense and respond to environmental cues through engineered genetic circuits^2–6^. A particularly compelling application lies in engineering autonomous sentinel plants—or phytosensors—that sense environmental threats and communicate them via remotely detectable, ideally quantifiable, outputs^7–10^. While certain plant species have long been used in agriculture as indicators of pests or diseases—and more recently for monitoring invasive plant pathogens in the context of climate change^11,12^—these traditional sentinel systems often suffer from low specificity, inconsistent performance, and a reliance on late-stage visible symptoms.

Recent advances in plant synthetic biology have enabled autonomous light emission in plants by expressing the fungal bioluminescence pathway (FBP) from *Neonothopanus nambi*, without the need for external substrates or excitation^13^. This pathway has since been repurposed as a versatile reporter for transcriptional circuit design^14,15^, with recent improvements including the successive enhancements of the pathway efficiency (eFBP)^16,17^, and the miniaturization of the system through the replacement of HispS with smaller, plant-derived alternatives^18^. These developments have opened new avenues for *in vivo*, substrate-free monitoring of plant processes at high spatiotemporal resolution^19,20^.

Here, we explore the potential of autobioluminescent plants as living biosensors for viral infection. We hypothesized that light-emitting genetic circuits could be deployed to create responsive sentinel systems, capable of translating viral presence into detectable light signals. Unlike luciferase or fluorescent reporters, autobioluminescence permits non-invasive imaging in real time using standard, low-cost optical equipment—making it particularly attractive for field-based phytodiagnostics.

We developed two mechanistically distinct strategies. In the first, we engineered recombinant plant viruses carrying a mini-HispS gene to complement transgenic *Nicotiana benthamiana* lines lacking this pathway component. Upon infection, the complete bioluminescent circuit was reconstituted, producing light specifically in infected tissues. In the second, we designed a virus-responsive Bioluminiscence Resonance Energy Transfer (BRET) circuit that integrates a luciferase–fluorescent protein fusion^21^ cleavable by NIa-Pro proteases encoded by members of the *Potyviridae*, the largest group of plant RNA viruses^22^. This dual-output system (Yellow-OFF/Green-ON) mimics electronic status indicators: uninfected plants emit yellow light via sustained BRET, while infection triggers a spectral shift towards green due to protease-mediated cleavage, providing a virus-specific optical signature.

Together, these strategies establish a modular framework for phytosensoring. By coupling synthetic bioluminescent circuits with viral sensing mechanisms, we demonstrate the feasibility of creating programmable, substrate-free sentinel plants capable of reporting infections remotely with specificity, autonomy, and spatiotemporal precision.

## Results

### Spatiotemporal viral tracking with autobioluminescence reveals the uncoupling between local and systemic responses

We first assessed whether luminescence could be used to monitor viral infections at the organ and organism level using minimal imaging equipment. For this purpose, FBPΔHispS *Nicotiana benthamiana* plants stably transformed with three of the four genes in the FBP (H3H, Luz and CPH), were agroinfected with viral vectors carrying either Hms, ASCL, or PKS2 genes, all three early described by Palkina et al ^18^ as small-size plant alternatives of the large fungal hispidin synthase gene. The viral systems assayed comprised: two agrobacterium infective clones of the tobacco mosaic virus (TMV) capable of cell-to-cell and systemic movement, and a second one deprived of systemic movement (TMVΔCP); a potato virus X (PVX) infective clone enabled for cell-to-cell and systemic movement; and a non-mobile deconstructed system based on the bean yellow dwarf virus (BeYDV)^23,24^ (Figure 1a).

**Figure 1.**
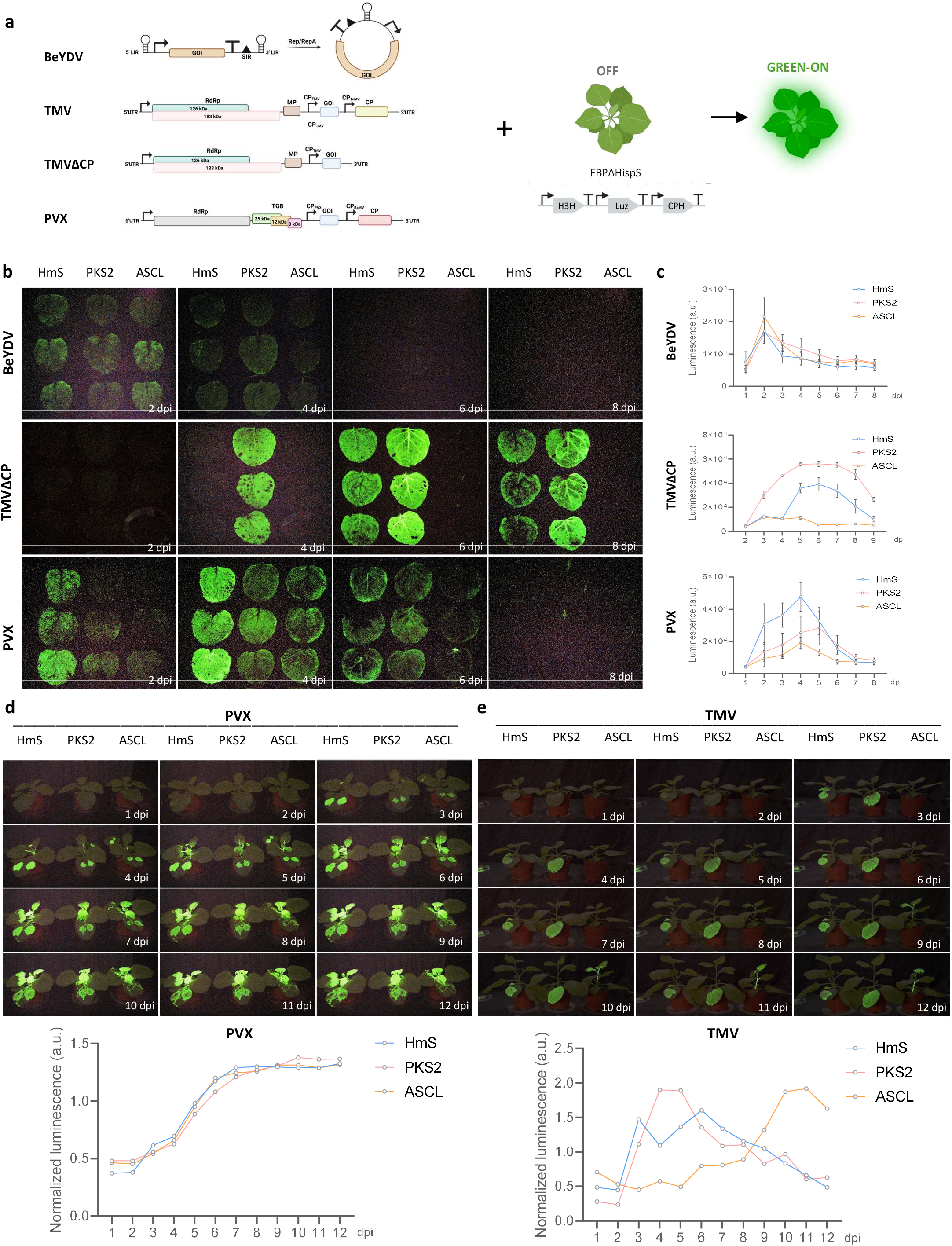
Autobioluminescence for viral tracking of BeYDV, TMV and PVX. **(a)** Schematic representation of the T-DNA constructs used and mode of action for the tracking of BeYDV, TMVΔCP, TMV and PVX viruses, including the stably transformed module for the constitutive expression of the FBPΔHispS genes (H3H, LUZ and CPH) and the BeYDV, TMVΔCP, TMV and PVX replicons carrying the different mini HispS substitutes (HmS, PKS2 or ASCL). **(b)** Time course of BeYDV (top), TMV (center) and PVX (bottom) local infection in FBPΔHispS *N*.*benthamiana* plants after infiltration of leaves expressing the different HispS substitutes (HmS, PKS2 and ASCL) and its quantification in **(c)**. Time course of PVX **(d)** and TMV **(e)** systemic infection in FBPΔHispS *N*.*benthamiana* plants after agroinoculation of basal leaves expressing the different mini HispS genes (HmS, PKS2 and ASCL).

Virus-associated luminescence levels were assessed in darkness using consumer-grade Digital Single-Lens Reflex (DSLR) cameras optimized for low-light detection linked to a customized lab management platform ^25^. A dedicated software tool was developed to quantify luminescent signals within defined regions of the captured images. To monitor organ-level dynamics, fully agroinfected leaves were detached post-infiltration, placed in Petri dishes under short-day photoperiod conditions, and imaged every 24 hours. Under these settings, marked differences in luminescence kinetics were observed among the three viral systems tested (Figure 1b, quantified in Figure 1c). The BeYDV replicon reached peak luminescence at 3 dpi, whereas the PVX and TMVΔCP systems peaked three and four days later, respectively. All three mini-HispS variants were functional across the systems, with the exception of ASCL, which exhibited undetectable local activity when mobilized via a TMVΔCP infectious clone.

We next evaluated the ability to trace and quantify the systemic spread of luminescence following local agroinfection of basal leaves with TMV and PVX clones (Figure 1d, 1e). When delivered via a PVX vector, all three HispS substitutes induced detectable luminescence in systemic tissues by 4 dpi— just 24 hours after light emission first appeared in locally infected leaves (Fig 1d, upper panel). The infection dynamics were comparable across the three PVX clones, as illustrated by the time-course of normalized luminescence (Fig. 1d, lower panel). Notably, detailed inspection of systemic PVX-infected tissues revealed that luminescence consistently preceded the onset of visible symptoms (Supplementary Figure 1). As expected, Hms and PKS2—but not ASCL—triggered local luminescence when delivered via TMV (Figure 1e, upper panel). Remarkably, and in contrast to initial expectations, only the TMV-ASCL infectious clone induced detectable luminescence in systemic tissues. The time-course analysis of normalized luminescence highlights this divergent behaviour, clearly distinguishing ASCL from Hms and PKS2 (Figure 1e, lower panel). These results underscore a functional uncoupling between local and systemic responses, suggesting that strong local activation does not necessarily predict the establishment of systemic infection.

### Engineering of a BRET-based spectral switch via targeted proteolytic cleavage

Having established a method for detecting viral infections with minimal instrumentation, we next set out to engineer a potyvirus-responsive sentinel circuit (Figure 2a). Central to this design was the development of a bioluminescent colour switch based on BRET, using LUZ as the energy donor\ (emission range: 460–625 nm). Two fluorescent proteins, DsRed and PdaC1, were tested as potential BRET acceptors based on excitation spectrum compatibility (Figure 2b).

**Figure 2.**
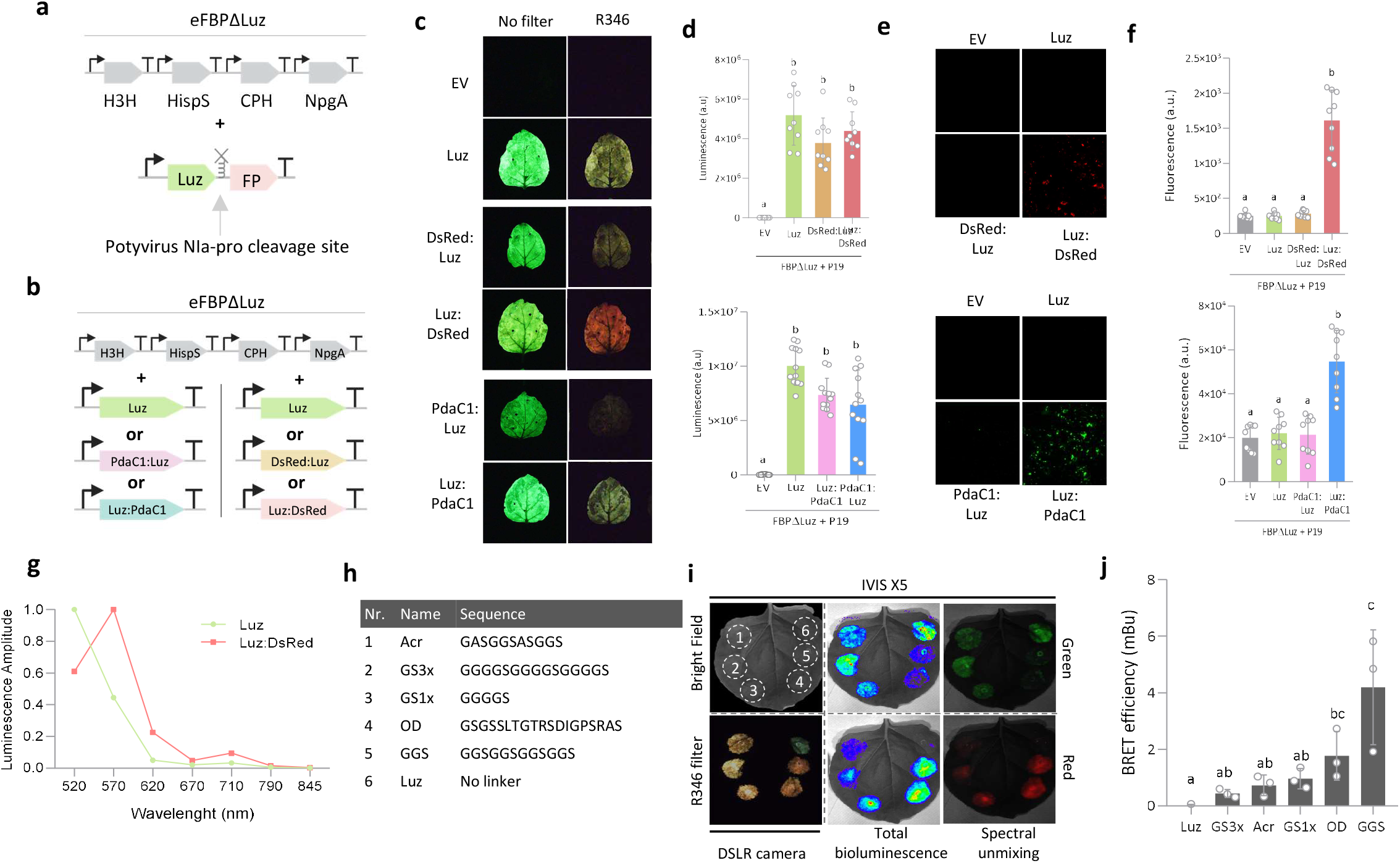
Design and characterization of the BRET switch. **(a)** Schematic representation of the T-DNA constructs used for the BRET-based Potyvirus sensor, including the module for the constitutive expression of the eFBPΔLUZ genes (H3H, HispS, CPH and NPGA) and the transcriptional unit of LUZ fused to a fluorescent protein (FP) with a linker carrying the Potyvirus NIa protease cleavage site. **(b)** Schematic representation of T-DNA constructs used for the BRET reporter system, with constitutive expression of the FBPΔLUZ genes (H3H, HispS, CPH and NPGA) and transcriptional units encoding LUZ fused to DsRed or PdaC1 in both N- and C-terminal configurations. **(c)** Bioluminiscence of leaf sectors agroinfiltrated with BRET constructs, imaged with a DSLR camera with and without a ROSCO R346 filter **(d)** Luminometer quantification of leaf disks taken from samples in (c). **(e)** Confocal fluorescence images and **(f)** fluorimeter-based quantification of the same samples. **(g)** Luminescence spectra deconvolution of LUZ and LUZ:DsRed. **(h)** Codenames and amino acid sequences of linkers tested in this study. **(i)** Imaging results of LUZ:DsRed fusions containing the linkers from (h), co-expressed with FBPΔLUZ. Images include bright field, green and red luminescence using the IVIS Spectrum X5 system, and bioluminescence using a DSLR camera and ROSCO R346 filter. Total bioluminescent signal detected using the red and green emission filters. Spectral unmixing was applied to separate overlapping emission spectra, allowing quantification of luminescence in the green and red channels. **(j)** Quantification of BRET efficiency for the LUZ:DsRed fusions generated with the linkers listed in (h), based on image analysis from (i). Statistical analyses were performed using One-way ANOVA (Tukey’s multiple comparisons test, P-Value ≤ 0.05). Error bars represent standard deviation (n = 9 for (d) and (f) and n=3 for (j)). Figure includes images from Biorender (biorender.com).

To evaluate energy transfer, we generated fusion constructs by placing each acceptor at either the N- or C-terminus of LUZ, connected via a flexible GGGGS linker (GS1X). These constructs were transiently expressed in *N. benthamiana* alongside the remaining components of the FBP lacking LUZ (eFBPΔLUZ) and visualized using a DSLR camera. Only the C-terminal fusions—particularly LUZ:DsRed—produced a visible shift in luminescence from green to a yellow colour (resulting from the combined red and green emissions caused by partial BRET efficiency), which was further enhanced in DSLR images by using a purple dichroic filter (Figure 2c).

BRET was accompanied by a modest, non-significant reduction in total luminescence (Figure 2d). Upon external excitation, DsRed fluorescence was observed exclusively in C-terminal fusions, as confirmed by confocal microscopy (Figure 2e) and fluorometric analysis (Figure 2f), and displayed a punctate pattern consistent with endomembrane association (Supplementary Figure 2). Based on the distinct and robust spectral shift, we selected the LUZ:DsRed fusion for further BRET optimization. Spectral emission profiles were confirmed using a charge-coupled device (CCD) camera (Figure 2g), and a series of alternative linker sequences were tested to improve energy transfer (Figure 2h). Among these, the so-called OD and GGS linkers yielded the highest BRET efficiency (Figures 2i, 2j) and were retained for subsequent experiments.

We next introduced an optimized tobacco etch virus (TEV) NIa-Pro protease cleavage site (CS) into the BRET linker to generate a colour switch responsive to site-specific proteolysis (Figure 3a). Insertion of the CS did not significantly affect BRET efficiency in any of the tested linkers (Supplementary Figure 4). All LUZ:linker-CS^TEV^:DsRed constructs were then co-infiltrated in *N. benthamiana* leaves with or without co-expression of the TEV NIa-Pro protease, following the experimental design outlined in Figure 3b. A shift in luminescence from yellow to green was observed exclusively in the presence of the protease, indicating successful cleavage and release of DsRed. This spectral switch was detectable both by filtered DSLR camera and with a CCD camera equipped with spectral analysis system (Figure 3c). Quantification of the 520/570 nm green-to-red (G/R) luminescence ratio confirmed the spectral shift, which was more pronounced for constructs carrying the OD and GGS linkers (Figure 3c).

**Figure 3.**
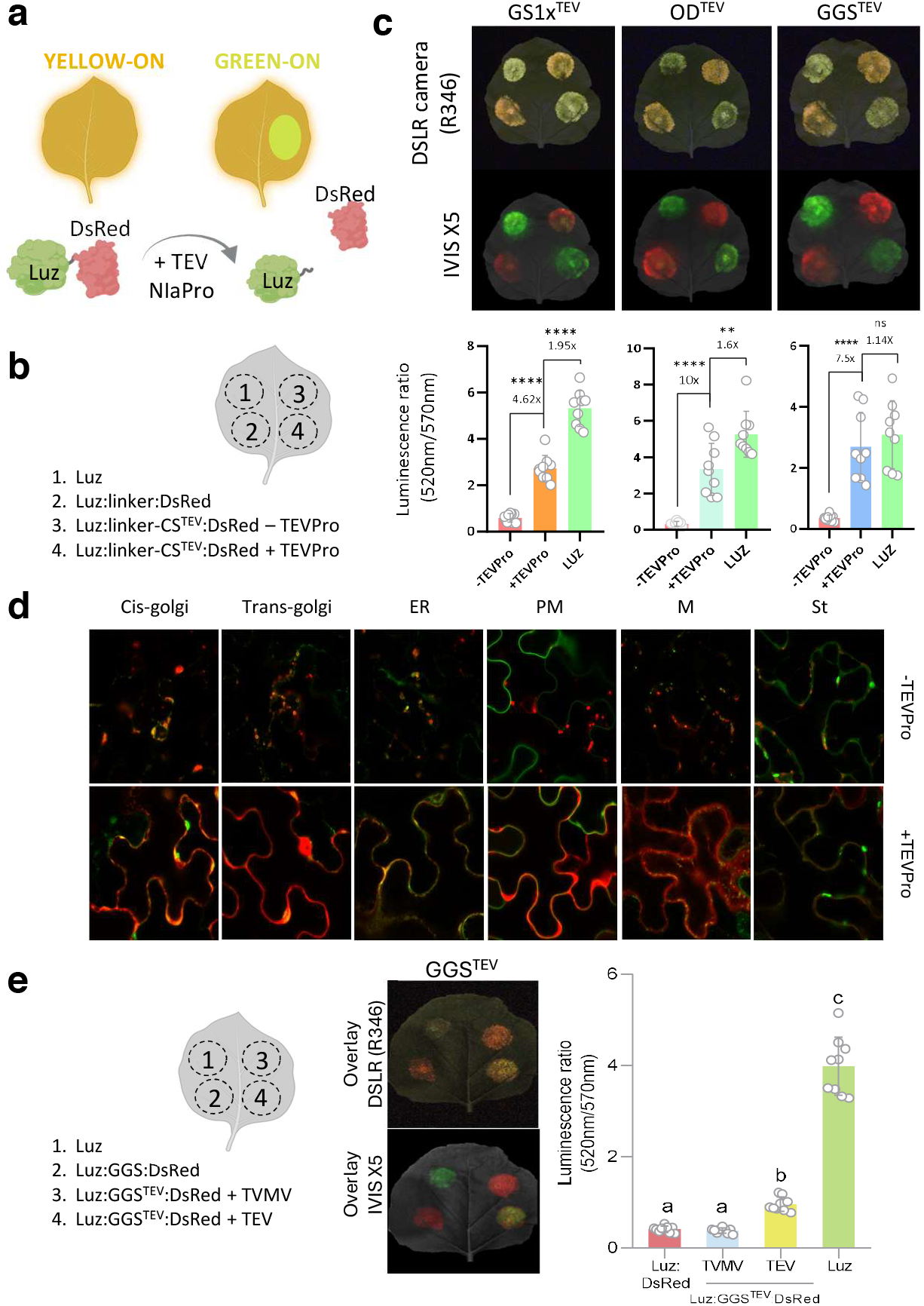
Implementation of a BRET-based Potyvirus switch. **(a)** Schematic representation of the mode of action of the BRET-based switch for Potyvirus detection. LUZ is fused to DsRed via a protease-sensitive linker containing the recognition site for the TEV NIa protease (TEVPro-(CS^TEV^)). Upon infection, the viral protease cleaves the linker, physically separating DsRed from LUZ and disrupting energy transfer. This cleavage event results in a BRET efficiency drop and a measurable luminescence shift from yellow (DsRed plus LUZ emission) to green (direct LUZ emission). **(b)** Infiltration schema of the leaves shown in (c). **(c)** Top: Dark field images (R346 ROSCSO camera filter) and spectral unmixing overlays acquired with the IVIS X5 system. Images represent merged unmixed channels corresponding to green and red emission components. Bottome: luminescence ratios (520nm/570nm) calculated from the IVIS X5 images. **(d)** Confocal microscopy characterization of *N. benthamiana* leaves infiltrated with Luz:GGS^TEV^:DsRed without or with TEVPro, co-infiltrated with a panel of subcellular markers targeting the cis- and trans-Golgi, endoplasmic reticulum (ER), plasma membrane (PM), mitochondria (M), and chloroplast stroma (St), as well as the eFBPΔLUZ module and P19. **(e)** Left: infiltration scheme of the depicted leaf. Middle: Dark field image (R346 ROSCSO camera filter) and overlay of spectrally unmixed channels obtained with the IVIS X5 system. Right: Luminescence ratios (520nm/570nm) in *N*.*benthamiana* leaves infiltrated with the Luz:GGS^TEV^:DsRed switch in the presence of TVMV or TEV, the eFBPΔLUZ module and P19. Statistical analyses were performed using One-way ANOVA (Tukey’s multiple comparisons test, P-Value ≤ 0.05). Error bars represent standard deviation (n = 9). Figure includes images from Biorender (biorender.com).

To investigate the spatial dynamics of the cleavage event, the subcellular localization of DsRed was examined by confocal microscopy (Figure 3d). In the absence of the protease, DsRed localized to endomembrane-associated compartments, as confirmed by colocalization with green fluorescent organelle markers. Upon NIa-Pro co-expression, DsRed signal redistributed to the nucleo-cytoplasm, consistent with protease-mediated release of the fluorescent moiety.

To further validate the functionality of the spectral switch, we replaced the recombinant protease with an infectious viral clone. To assess specificity, we also included an infectious clone of the tobacco vein mottling virus (TVMV) in the assay (see Supplementary Figure 5). As shown in Figure 3e, agroinfiltration with the TEV infectious clone, but not with TVMV, induced a statistically significant shift in the 520/570 nm luminescence ratio in plants expressing the LUZ:GGS-CS^TEV^:DsRed sentinel construct. This colour change was readily detectable using both a CCD and DSLR camera systems (Figure 3e). Conversely, the TVMV-responsive construct displayed equivalent specificity, responding only to TVMV and not to TEV (Supplementary Figure 5).

### Genomic integration of the BRET-based spectral switch enables autonomous detection of TEV infection in *Nicotiana benthamiana*

To establish a *bona fide* glowing sentinel (Figure 4a), we generated two transgenic *Nicotiana benthamiana* lines co-expressing (i) the output luminescence module eFBPΔLUZ expressing HispS, H3H, CPH and NPGA and (ii) a BRET-based switch module, incorporating either the Luz:GGS-CS^TEV^:DsRed, or the Luz:OD-CS^TEV^:DsRed construct (GGS^TEV^ or OD^TEV^ lines, respectively). The functionality and specificity of the switch were then tested by transient expression of the TEV NIa-Pro protease by agroinfection with TEV, using TVMV as a negative control (Figure 4b). The OD^TEV^#15 line exhibiting the expected yellow background luminescence—indicative of an intact, non-cleaved OFF state—and the strongest green luminescent upon agroinfection was selected for further characterization.

Next, we assessed activation of the green-ON state in systemic tissues following agroinfection of basal leaves with the viral construct. In ODTEV#15 sentinel plants, this systemic response was first confirmed by spectral analysis of distal tissues using CCD camera imaging. As shown (Figure 4c), distal tissues exhibited a green-shifted yellow luminescence exclusively upon infection with TEV, but not with TVMV, confirmed by quantification of the G/R ratio in each plant (Figure 4d). The presence or absence of each virus in systemic tissues was validated by RT-PCR (Figure 4d). To evaluate sentinel performance using low-cost equipment, luminescence was then monitored over time in TEV- or TVMV-agroinfected OD^TEV^#15 plants using an unfiltered DSLR camera (Figure 4e). Under these conditions, a faint yellow glow—corresponding to the yellow-OFF state—was visible from the start of the experiment, primarily in the stem, and was independent of infection status. Upon TEV infection, a green-ON luminescent signal first appeared in locally inoculated leaves at 3 dpi, followed several days later by robust green luminescence in apical tissues. The signal then propagated basipetally, following a stereotypical vasculature-first spread pattern. This dynamic produced a characteristic time-course profile that was both data-rich and readily captured using simple luminescence measurements (Figure 4f).

**Figure 4.**
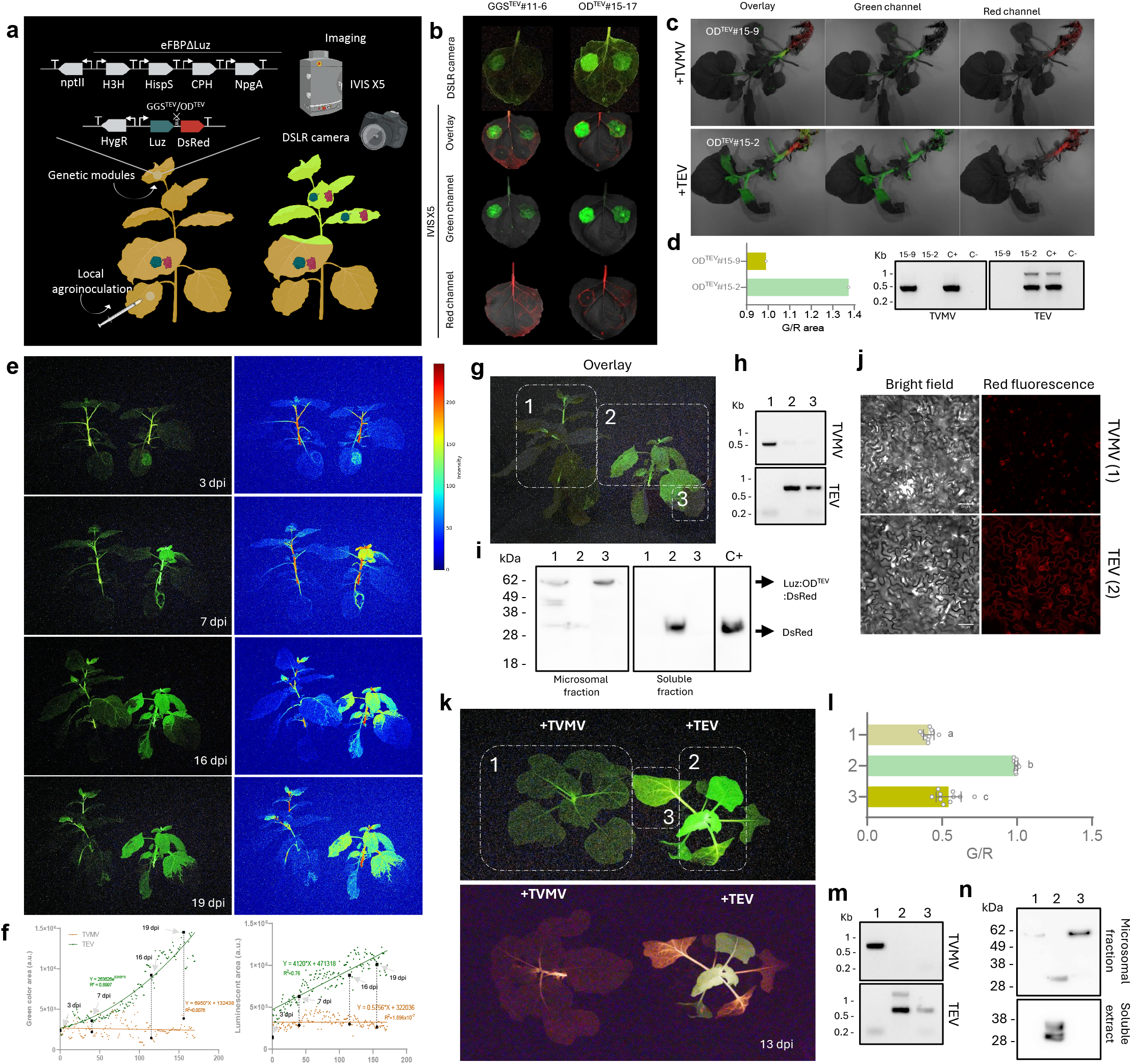
BRET-based viral sentinels for real-time detection of TEV protease activity in plants. **(a)** Schematic representation of the genetic constructs used to generate the TEV sentinels and their action mechanism. **(b)** Bioluminescence images acquired with a DSLR camera and the IVIS X5 system showing GGS^TEV^ and OD^TEV^ T1 leaves infiltrated with TEV NIaPro (top left), TEV (top right), TVMV NIaPro (bottom left) and TVMV (bottom right). **(c)** Representative IVIS X5 overlays of unmixed green and red channels in whole T1 plants agroinoculated with TVMV (top) and TEV (bottom) at 6 dpi. **(d)** Left: green-to-red ratio (G/R) obtained by quantification of the pictures in (c). Right: RT-PCR of appex (A) and root (R) samples of ODTEV15-2 (2) and OD^TEV^15-9 (9) confirming the infection of T1 plants shown in (c). **(e)** Time-course bioluminescence images acquired with a DSLR camera (left) and color maps (right) showing the temporal dynamics of luminescence signal in plants infected with TVMV (left) or TEV (right). **(f)** Quantification of the total green area (left) and luminescence intensity (right) over time for the plants shown in (e). **(g)** Bioluminescence image of plants from (e) at the end of the experiment indicating samples collected for molecular analyses. (1) TVMV agroinoculated plant; (2) luminescent and (3) non-luminescent tissue from the TEV agroinoculated plant. **(h)** RT-PCRs of the plants shown in (g) confirming infection. **(i)** Anti-DsRed western blot detection confirming Luz:OD^TEV^-DsRed cleavage in planta upon TEV infection carried out with the same samples from (g). Expected sizes: Luz:OD^TEV^-DsRed (58.8 KDa), DsRed (27.5 KDa) **(j)** Confocal microscopy of TVMV and TEV infected plants. **(k)** Top view bioluminescence image acquire with a DSLR camera without (top) and with R346 filter (bottom) of T2 OD-TEV#15 plants agroinfected with TVMV and TEV at 13 dpi. (1) TVMV agroinoculated plant; (2) green-luminescent and (3) red-luminescent tissue from the TEV agroinoculated plant. **(l)** Green-to-red ratio (G/R) of leaves from (1), (2), and (3) obtained by quantification of picture in k (bottom). Error bars represent standard deviation (n = 10). Statistical analysis was performed using a one‐way ANOVA with Tukey’s multiple comparisons test (P‐value ≤0.05). Variables with the same letters belong to the same statistical group. **(m)** RT-PCRs of the plants shown in (k) confirming infection. **(n)** Anti-DsRed western blot detection confirming Luz:DsRed cleavage in planta upon TEV infection carried out with samples from (k). Figure includes images from Biorender (biorender.com).

The correct functioning of the sentinel was confirmed by analyzing selected tissue samples from TEV- and TVMV-infected plants (depicted in Figure 4g). The presence of each virus was first verified by RT-PCR (Figure 4h), followed by assessment of the subcellular distribution of the Luz:DsRed fusion partners. Notably, the systemic switch to the green-ON state correlated with the expected molecular and subcellular signatures, including the translocation of DsRed signal from microsomal to soluble fractions as detected by Western blot (Figure 4i) and a redistribution of red fluorescence from endomembrane-associated regions to nucleocytoplasmic compartments observed by confocal microscopy (Figure 4j).

In late-stage TEV infections, basal leaves occasionally displayed luminescence patterns that deviated from the typical vasculature-first spread (Figure 4k, upper panel), raising questions about their infection status. As shown in Figure 4k (lower panel), the yellow-OFF/green-ON dual-output logic— captured using a DSLR camera with a purple filter—enabled accurate discrimination of infection states, independent of absolute luminescence intensity. TVMV-infected tissues showed the lowest G/R ratios, while TEV-infected apical tissues showed the highest (Figure 4l). Basal leaves from TEV-infected plants exhibited significantly lower G/R ratios than apical tissues, consistent with low but RT-PCR-detectable viral loads and predominant microsomal localization of DsRed (Figure 4m, n).

To further evaluate sentinel specificity, we challenged plants with two phylogenetically distant viruses—PVX and TMV (Figure 5a). As with TVMV, neither PVX nor TMV triggered the green-ON state; only TEV induced the characteristic luminescence time-course associated with infection (Figure 5b).

**Figure 5.**
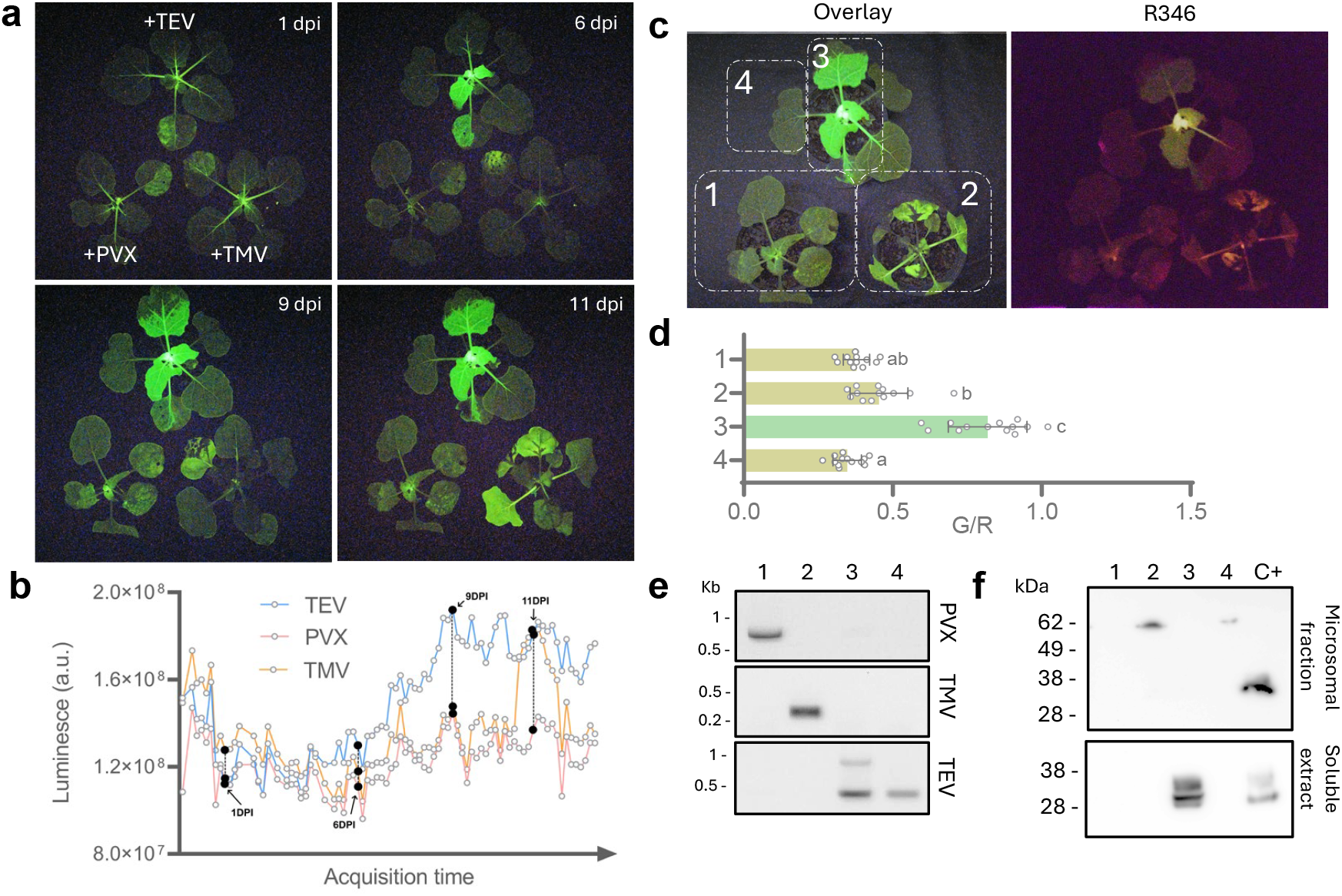
Specificity of BRET-based viral sentinels. **(a)** Time-course dark shot images acquired with a DSLR camera showing the temporal dynamics of luminescence signal in plants agroinfected with TEV (top), PVX (bottom left) or TMV (bottom right). **(b)** Quantification of the luminescence intensity over time for the plants shown in (a). **(c)** Dark shot and filter (R346) pictures taken with a DSLR camera at 12 dpi. (1) PVX-(2) TMV-agroinoculated plants; (3) green-luminescent and (4) yellow-luminescent tissue from a TEV agroinoculated plant. **(d)** Green-to-red ratio (G/R) obtained by quantification of the picture taken with the R346 filter and displayed in (c). Error bars represent standard deviation (n = 12). Statistical analysis was performed using a one‐way ANOVA with Tukey’s multiple comparisons test (P‐value ≤0.05). Variables with the same letters belong to the same statistical group. **(e)** RT-PCRs confirming the infection for the PVX-, TMV- and TEV-infected plants shown in (b). **(f)** Western blot detection confirming LUZ:DsRed cleavage in planta upon TEV infection carried out with samples from (b). Expected sizes: Luz:OD^TEV^-DsRed (58.8 KDa), DsRed (27.5 KDa).

As previously observed in late-stage TEV infections, elevated total luminescence occasionally appeared in necrotic basal leaves during advanced TMV infection (Figure 5c). However, filtered DSLR images clearly distinguished infection states, revealing a yellow-OFF output (low G/R ratios) in TMV-infected basal leaves, in contrast to the green-ON signature (high G/R ratios) in apical leaves of TEV-infected plants (Figure 5d). Molecular diagnostics confirmed the infection status of all samples (Figures 5e and 5f). These findings underscore the robustness of the dual-colour system in preventing false positives under complex pathological scenarios.

## Discussion

Early and reliable detection of crop pathogens is increasingly critical for sustainable agriculture, particularly under the pressures of climate change and reduced pesticide use. Yet conventional diagnostic methods such as PCR and ELISA—though accurate—are invasive, offline, and reliant on laboratory infrastructure, limiting their scalability for in-field use. Passive spectral techniques (e.g., NIR, thermal imaging) offer non-invasive alternatives but typically rely on indirect physiological signatures and complex interpretation models. Biosensors and nanomaterials have been proposed to monitor plant health by detecting metabolites such as H_2_O_2_ and the phytohormone salicylate^26,27^, which are known to accumulate at high levels during biotic stress, including viral infections. However, physiological or metabolic signatures may not be reliable indicators of pathogen presence, as some infections can remain asymptomatic or latent^28^. For example, negative autoregulation in potyviruses has been reported as an immune evasion strategy that modulates infection kinetics to avoid the activation of phytohormone signaling pathways^29^.

Active phytosensing systems report defined molecular events, thus offering greater specificity and interpretability. Our work addresses this need by establishing a robust and modular platform for autonomous pathogen detection in plants, integrating synthetic signal processing with visible optical outputs. By embedding a protease-responsive BRET circuit into a bioluminescent chassis, we engineered *Nicotiana benthamiana* plants that act as living sentinels, capable of reporting viral infections through both spectral and intensity-based cues. This represents one of the most advanced real-world applications of plant synthetic biology to date, and a tangible step toward field-deployable sentinel plants for precision crop protection.

The system’s core innovation lies in its ability to detect virus-specific proteolytic activity *in vivo*. The TEV-responsive switch reliably distinguishes infected from non-infected plants and even resolves infection boundaries within a single plant. Remarkably, this detection is achieved using low-cost imaging tools, highlighting the potential for real-time, remote monitoring without external reagents or invasive sampling. A key conceptual advance is the implementation of a dual-output design that mimics status-indicator logic in electronic alarm systems: not only does the system report infection through a spectral switch, but it also provides a stable signal in healthy plants, confirming system integrity. This “pilot light” feature is critical to minimize false negatives and ensure confidence in the absence of infection—an often-overlooked but essential attribute for practical deployment.

Although initially developed for TEV and extended to TVMV, the approach is broadly generalizable. Numerous plant pathogens—including viruses, bacteria, and fungus—deploy proteases or intracellular effectors with defined cleavage specificities. BRET linker peptides can be redesigned for specificity toward the over 200 known potyviruses^22^, including potato virus Y (PVY), one of the most economically damaging plant viruses worldwide^30^. Conversely, NIa-Pro cleavage motifs themselves can be engineered to tune the switch’s sensitivity, ranging from highly specific recognition—as demonstrated here and by others^31^—to broader-spectrum responsiveness, as recently shown by Wang et al.^32^. Moreover, bacterial effectors such as AvrRpt2^33^ and AvrPphB^34^ offer opportunities for tailored biosensing beyond viral targets. These effectors cleave host proteins at well-defined motifs, which can be repurposed in synthetic circuits to detect pathogen activity^35^. Together, these features position protease-activated switches as a versatile and modular platform for real-time plant health surveillance across a broad spectrum of pathogen types.

In our experiments we observed that protease activation led not only to a spectral shift but to a modest increase in luminescence intensity. This may reflect improved enzymatic performance upon cleavage of the LUZ:DsRed fusion—possibly due to changes in subcellular localization, or relief of steric hindrance or quenching. Alternatively, the activation of defense mechanisms and/or necrotic reactions could enhance flux through the phenylpropanoid pathway, increasing availability of caffeic acid, the FBP substrate. This later mechanism would also explain the increase in yellow luminescence observed during late stages of both specific and non-specific infections. Either mechanism underscores the sensitivity of bioluminescence to intracellular context and stress physiology.

Future developments may focus on increasing the brightness and versatility of the system. Enhanced luciferase variants, BRET acceptors with higher quantum yield, and pathways emitting at shorter wavelengths—such as the five-colour Nano-lanternX (NLX) system^36^—to boost signal strength and enable expanded multiplexing. Integration of orthogonal sensors into multicolours sentinel lines offers a promising route toward simultaneous detection of diverse pathogens or environmental cues. While *N. benthamiana* is a particularly convenient host for viral detection due to its broad susceptibility^37^, its utility as a dedicated phytosensor chassis could be further improved through targeted engineering—such as suppressing immune responses to enhance sensitivity and maintain signal integrity^38^. Altogether, this work provides a proof-of-concept for whole-plant biosensors capable of decoding pathogen-derived molecular signals into visible readouts. By leveraging synthetic biology to make infections visible, bioluminescent sentinels may help shift plant disease management from reactive to predictive strategies, with profound implications for sustainable agriculture.

## Materials and methods

### Modular cloning

DNA constructs used in this work were assembled using GoldenBraid (GB) modular cloning system^41^. Coding sequences of proteins used in this work were codon optimized for *Nicotiana benthamiana* using IDT Codon Optimization Tool (https://eu.idtdna.com/CodonOpt) and subsequently domesticated using the GB Domesticator tool (https://goldenbraidpro.com/do/synthesis/). Level 0, Level 1 and Level>1 assemblies were performed using GB protocol as previously described by Sarrion-Perdigones et al.^41^. Level 0 parts were confirmed by restriction analyses and Sanger sequencing, and Level 1 and Level >1 assemblies were verified by restriction analyses. All plasmids used in this work are listed in Table S1.

### N. benthamiana transient expression

Plasmids were transferred to *Agrobacterium tumefaciens* strain GV3101 or EHA105 (see Table S1 for details). Five-to-six week-old *N. benthamiana* plants grown under controlled conditions (24°C,16 h light/ 20°C, 8 h dark) were used for agroinfiltration. Overnight *A. tumefaciens* cultures were pelleted and resuspended in infiltration buffer (10 mM MES, pH 5.6; 10 mM MgCl_2_; 200 µM acetosyringone) to an optical density of 0.1 at 600 nm (OD_600_). Bacterial suspensions were incubated for 2 hours at room temperature on a horizontal rolling mixer in darkness. For co-expression experiments different *Agrobacterium* cultures harboring different GB elements were mixed prior to infiltration. Finally, infiltration was performed using a 1-ml needleless syringe, applied to the abaxial side of the three youngest fully expanded leaves per plant. Detailed information on the experimental design can be found in the Supporting Information section.

### N. benthamiana viral agroinoculation

Binary vectors of BeYDV, PVX, TMV, TEV and TVMV clones were reported^23,42–45^. Plasmids were transferred to the *A. tumefaciens* strains EHA105 or GV3101. Single colonies were grown at 28°C for 24 hours, pelleted and resuspended in agroinfiltration buffer to an OD_600_ of 0.1 for PVX, TMV and BeYDV, and OD_600_ of 0.3 for TEV and TVMV. Agroinfections for BeYDV, TMV and PVX tracking were performed in *N. benthamiana* transgenic lines constitutively expressing three out of the four genes of the fungal bioluminescence pathway (CPH, H3H and LUZ) under CaMV 35S promoter. For local infection assays, infiltrated leaves were detached at 1 day post infiltration (dpi) for PVX and BeYDV, and at 2 dpi for TMV, and placed on Petri dishes containing water. For systemic infection assays with PVX, TMV, TEV and TVMV, one or two basal leaves per plant were agroinoculated. Plants and leaves were maintained under 16 h light / 8 h dark photoperiod and photographed every 2 or 24 hours, depending on the experiment.

### RT-PCR analyses

Total RNA was extracted from meristem and systemic leaves for viral RNA detection using GeneJEt Plant RNA Purification Kit (Thermo Scientific). First-strand cDNA was synthesized from 1 ug of total RNA using PrimeScript RT Reagent Kit (Takara Bio), according to manufacturer’s instructions. Specific primers for each virus (listed in Table S2) and MyTaq DNA polymerase (Bioline, Meridian Bioscience) were used, following manufacturer’s protocol.

### Image acquisition and processing

Bioluminescence pictures were taken in a dark room with a Sony Alpha ILCE-7M3 camera (DSLR camera) with a Sony SEL20F18G objective (ISO 51200, f/2.0 and 30 seconds of exposure). Images were acquired in uncompressed RAW (ARW) format and processed using Sony Imaging Edge Desktop software to convert them to 8-bit TIFF images for visualization and 16-bit TIFF images for quantification. Color separation for BRET-based experiments was achieved by using Permacolor dichroic ROSCO filters: R343, R346, R349 and R49. The R343, R346, and R349 filters transmit approximately 33%, 22%, and 25–30% of incident light, respectively, corresponding to light losses of –1.6 to –2.2 f-stops. The R49 filter transmits about 15% (occasionally estimated as low as 25%), resulting in a loss of –2.5 to –3.0 f-stops. All filters allow significant transmission in the violet (380– 450 nm) and deep red (600–700 nm) regions, while strongly attenuating green to yellow wavelengths (500–580 nm). Filter details are available on ROSCO’s MyColor Selector: https://emea.rosco.com/es/node/1706. Luminescence pictures of stable plants with R346 filter displayed in Figures 4 and 5 were taken with 5 min of exposure time (ISO 51200, f/1.8).

### Luminescence and fluorescence quantification

For the luminescence quantification of full leaves and plants displayed in Figure 1, images were processed using Fiji software. Green channel was selected for bioluminescence quantification. Total plant tissue was defined as region of interest (ROI) for total intensity measurements, and area selected was used for normalization. Mean values are reported.

For luminescence and fluorescence measurements from leaf discs (Figure 2), they were collected at 72 hours post infiltration using a cork borer with a diameter of 0.5 cm. Two or three discs per agroinfiltrated leaf were excised. Leaf discs were transferred to a black 96-well microplate. Bioluminescence measurements were carried out with a GloMax^®^-Multi Detection System (Promega) by using an integration time of 10s. DsRed2 (Ex λ _max_: 561 and Em λ _max_: 587) and PdaC1 (Ex λ _max_: 480 and Em λ _max_: 492) fluorescence activities were determined with Perkin Elmer Victor X2 Microplate Reader.

For the quantification of luminescence in plants depicted in Figure 4, image analysis was performed using FIJI. The green (G) and red (R) channels were chosen for bioluminescence quantification. The total plant was designated as the ROI for measuring total and green color intensities displayed in Figure 4f. For Figure 4g, G/R ratios were calculated as total area values of the green channel against the red channel area values. For Figures 4l and 5d, 10-12 squares of 10^*^10 pixels were randomly selected from the indicated areas, and designated as the ROI for measuring intensity in the G and R channels. Average and individual values are shown.

### Confocal imaging

For subcellular localization experiments displayed in Figure 3 and Supplementary Figure 2, leaf samples were collected at 72 hours post infiltration. Fluorescence images were acquired using a ZEISS LMS 780 AxioObserver Z1 confocal laser scanning microscope equipped with a C Apo 40×/1.2 W water-immersion objective. Confocal images displayed in Figure 4 were obtained using a confocal Stellaris 8 FALCON (Leica) microscope. Fluorescence images from ZEISS 780 AxioObserver Z1 confocal microscope were analysed using ZEN 2.5 lite and ImageJ (FIJI) software and images from Stellaris 8 FALCON (Leica) with ImageJ (FIJI) software.

### BRET efficiency calculation

Leaf samples were collected at 72 hours post infiltration. Image acquisition and spectral unmixing was done in an IVIS Lumina-X5 Imaging System (Revvity) following manufacturer indications. Luminescence at 520nm (donor channel) and 570nm (acceptor channel) were used to determine BRET efficiency. BRET ratio was calculated using the equation described by Dimri *et al* ^46^. All luminescence images from IVIS X5 Lumina Imaging System were analysed using Revvity’s Living Imaging software and ImageJ (FIJI). For Revvity’s Living Imaging software, the automatic spectral unmixing tool was employed to decompose and process both luminescence (520-570nm) channels. From the resulting images, three regions of interest (ROIs) of equal dimensions were defined using FIJI and used to quantify the total counts per replicate per channel for subsequent data analysis.

### Computational analysis of viral bioluminescence

For the automated quantification of viral bioluminescence kinetics, a pipeline was implemented in Python 3.x with a graphical interface (PyQt5) that integrates OpenCV for image reading and pre-processing, NumPy for array handling, and Matplotlib for graph and colormap generation. The system was capable of generating timecourses of bioluminescence images with or without overlay with brightfield images, generating color maps for plants using a standardized intensity scale where minimum and maximum values were calculated globally from all regions of interest across all timecourse images to ensure visual consistency between frames, and performing comparative analysis through the extraction of absolute luminescent intensity values in predefined regions of interest. From 16-bit TIFF files (see Image acquisition and processing section), each bioluminescence frame was automatically converted to 8-bit grayscale (0-255) by applying the standard luminance formula (Y = 0.299 R + 0.587 G + 0.114 B) with R, G, and B taken in their original 16-bit range and internally scaled to the 8-bit range. Regions of interest (ROIs) are defined as rectangles of constant dimensions that are configured directly in the interface, and are extracted from each image by cropping the NumPy array according to their spatial boundaries. For quantification, in each ROI and frame, the values of all pixels in grayscale are summed, providing an absolute value of luminosity. Both processes utilized pixel value extraction from grayscale images in predefined regions of interest. To ensure visual comparability between all frames, the minimum (vmin) and maximum (vmax) intensity values are calculated globally considering all pixels from the selected ROIs throughout the entire time-course. From these data, the system automatically generates temporal series of luminescent intensity, timecourses, for each ROI. For the spatial representation of bioluminescence, the ‘jet’ colormap from Matplotlib is used, scaled with the global vmin/vmax values. Finally, all quantitative information is exported in CSV format with the corresponding luminescent intensity values for each analyzed ROI, and colormaps in PNG format, fully automated from the GUI. For comparison between viral treatments displayed in Figure 5 (TEV, PVX and TMV), comparative graphs were generated representing the temporal evolution of luminescent intensity for each virus.

### N. benthamiana stable transformation and selection of sentinel lines

*Agrobacterium tumefaciens* strain LBA4404 was co-transformed with two compatible plasmids carrying HygR-LUZ:OD^TEV^/GGS^TEV^:DsRed and nptII-FBPΔLuz ^47^ (see Table S1). *A. tumefaciens* culture was grown in MGL medium to exponential phase (OD_600_ 0.2-0.5) with plasmid-selective antibiotics and acetosyringone. Cells were then pelleted and resuspended in TY medium. Discs from five-to six-week-old *N. benthamiana* full-expanded leaves were cut and placed onto co-cultivation medium (MS medium, pH 5.7, supplemented with 1 mg/L 6-benzylaminopurine and 0.1 mg/L naphthalene acetic acid) for 24 hours. Discs were then immersed in the *Agrobacterium* suspension for 15 minutes and returned to co-cultivation medium for 48 hours. Then, discs were transferred to selection medium (containing 100 mg/L kanamycin, 20 mg/L hygromycin and 200 mg/L carbenicillin) for shoot regeneration. Regenerated shoots were transferred to rooting medium (hormone-free) for root development. Rooted plantlets were transplanted to soil and growth in a greenhouse under 16h light/8h dark photoperiod at 24ºC. A total of 19 GGS^TEV^ and 25 OD^TEV^ T0 plants were recovered. Among these, three GGS^TEV^ and three OD^TEV^ lines exhibiting strong luminescence throughout the entire plant were selected and self-pollinated to obtain the T1 generation. T1 seeds were germinated in MS selective media containing kanamycin (100 mg/L) and hygromycin (20 mg/L) and screened for luminescence at the seedling stage. Luminescence was detected only in two GGS^TEV^ (GGS^TEV^#5 and GGS^TEV^11) and one OD^TEV^ (OD^TEV^#15) lines. However, after two weeks, approximately half of the GGS^TEV^#5 seedlings lost luminescence, and this line was discarded for further experiments. Silencing was also observed in some T1 and T2 OD^TEV^#15 plants grown on selective media. Silenced segregants were discarded and only luminescent plants were used for subsequent experiments.

### Protein Extraction and Microsomal Fractionation

Collected plant tissue was flash-frozen in liquid nitrogen, ground to a fine powder, and stored at – 80 °C until use. For soluble protein extraction, tissue was homogenized in a 1:2 (w:v) ratio using phosphate-buffered saline (PBS; 137 mM NaCl, 2.7 mM KCl, 10 mM Na_2_HPO_4_, 1.8 mM KH_2_PO_4_, pH 7.4). The homogenate was centrifuged at 12,000 × g for 10 min at 4 °C, and the supernatant was collected as the soluble protein fraction. For microsomal fractionation, 200 mg of frozen tissue were processed according to the protocol described by Abas and Luschnig (2010)^48^, which involves differential centrifugation and phase partitioning using an aqueous polymer two-phase system to enrich for membrane-associated proteins. Pellets were resuspended in SDS sample buffer and incubated at 95 ºC for 5 min prior to analysis.

### SDS-PAGE and Western blot analysis

Proteins were separated by SDS-PAGE using MES-SDS running buffer (pH 7.3) on NuPAGE 10% Bis-Tris gels (Invitrogen) under reducing conditions. Then, proteins were transferred to a polyvinylidine difluoride (PVDF) membrane (Amersham Hybond™-P, GE healthcare) using a semi-wet blotting system (XCell II™ Blot Module, Invitrogen). Membranes were blocked with 2% ECL Prime blocking reagent (GE Healthcare) in PBS containing 0.1% (v/v) Tween-20 for 30 minutes with agitation at room temperature. For Ds-Red detection, membranes were incubated overnight at 4ºC with 1:500 dilution of anti-dsRed primary antibody (OriGene Technologies, Cat# TA180084) on a shaker, followed by incubation with 1:10000 dilution of HRP-labelled anti-mouse IgG secondary antibody (GE Healthcare, Cat# NA931) for 1 hour at room temperature. Signal detection was performed using ECL Prime Western blotting detection reagent (GE Heathcare) and visualized using a Quant500 imager for chemiluminescent detection with exposure times ranging from 1 min (for soluble fraction detection) to 30 min (for microsomal fraction detection).

### Sample size and statistical analysis

The sample number and statistical analysis considerations for each experiment are indicated in the corresponding figure. Statistical analyses were performed using One-way ANOVA (Tukey’s multiple comparisons test, P-Value ≤ 0.05). Plotting and statistic tests were all performed with GraphPad.

## Acknowledgements

The authors would like to thank José Antonio Navarro (IBMCP-CSIC) for providing the *A*.*tumefaciens* strains with the subcellular selection markers, Marisol Gascón (IBMCP-CSIC) for her assistance with confocal imaging acquisition and Alexander S Mishin, Andrey Gorokhovatsky and Tanya Chepurykh for valuable discussions on Luz:DsRed detection.

