## Supplementary information for "Bioluminescent sentinel plants enable autonomous diagnostics of viral infections"

### Supporting Information

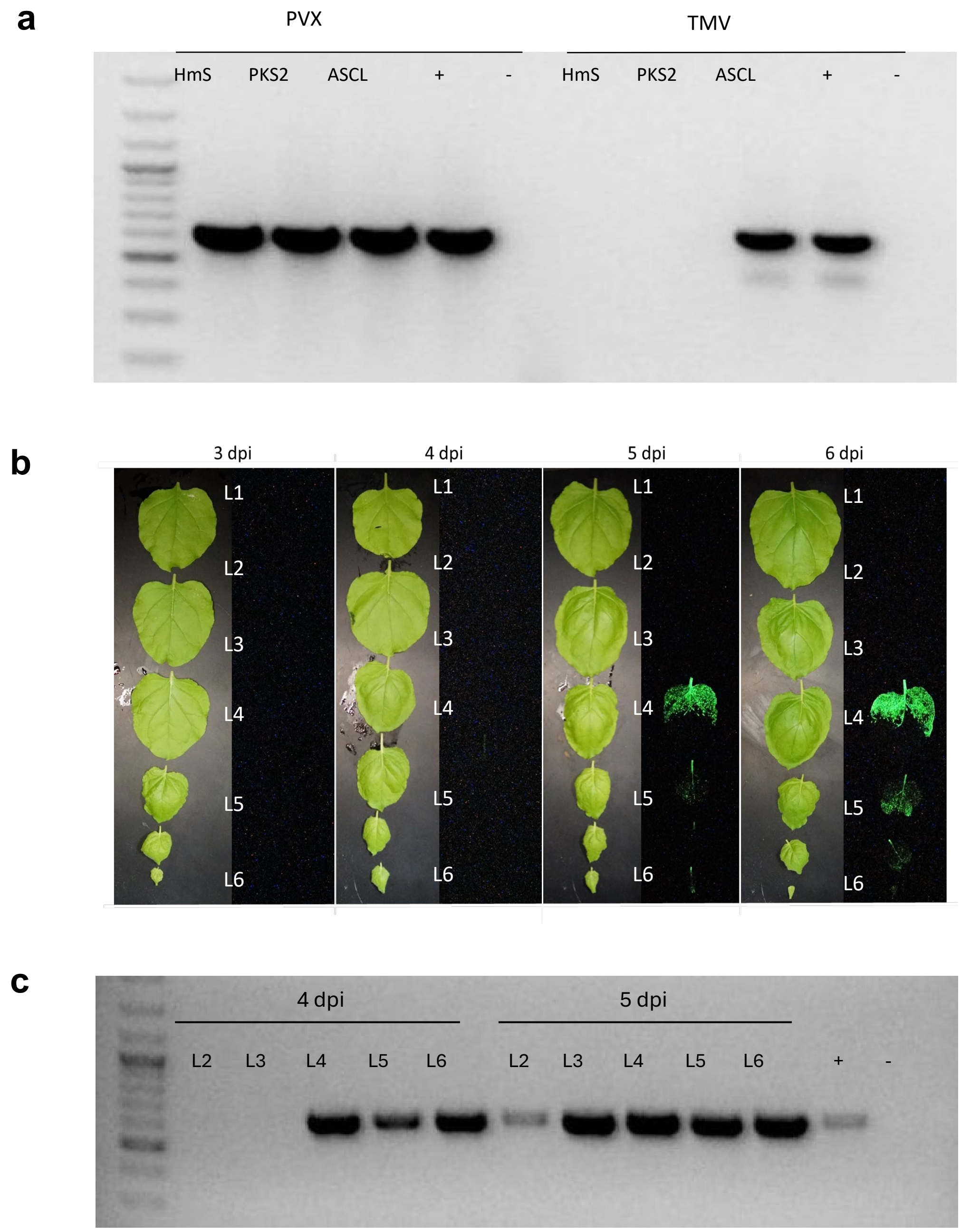


**Supplementary Figure 1.** (a) RT-PCR analysis of systemic tissue from plants of Figure 1d infected with PVX and TMV expressing the three Hispidin synthase susbtitutes. (b) Systemic leaves from FBPΔHispS *N. benthamiana* plants agroinoculated in basal leaves with PVX:HmS, detached for symptoms observation and bioluminescence detection comparison. (c) RT-PCR analysis for PVX detection in selected detached leaves, both bioluminiscent and non-bioluminiscent.

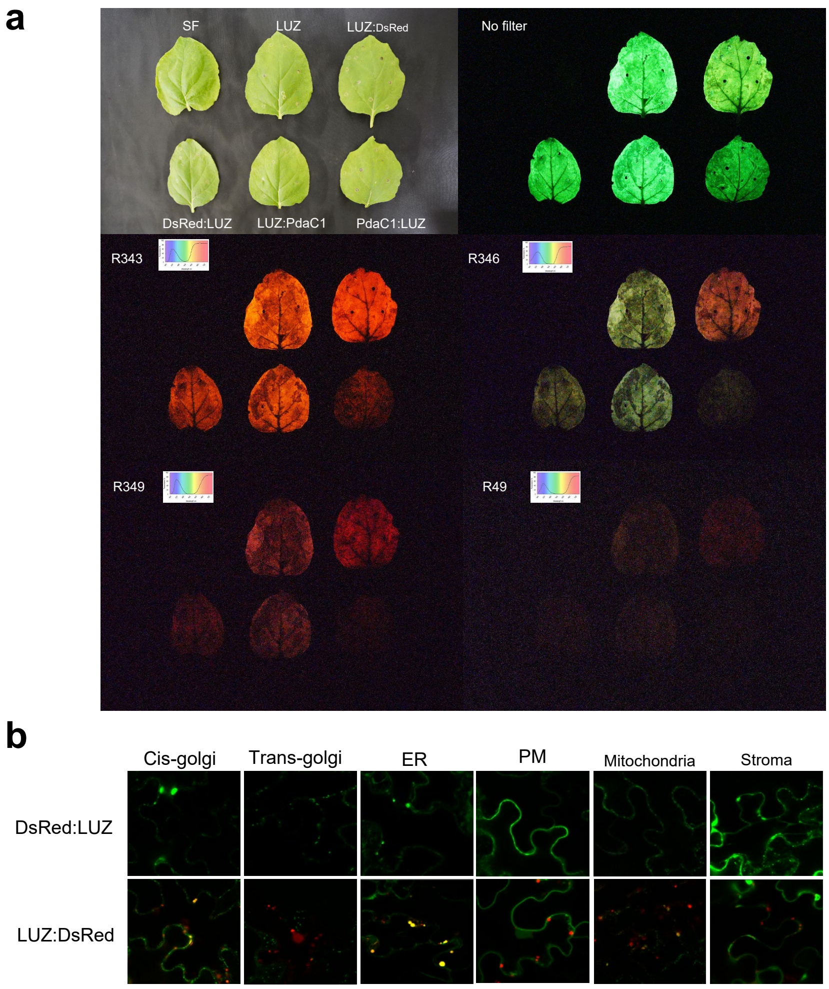


**Supplementary Figure 2.** (a) Bright and Dark field (taken with ROSCO's R343, R346, R349 and R49 filters) camera acquisitions of *N. benthamiana* leaves infitrlated with LUZ, LUZ-DsRed or LUZ-PdaC1 (in both N- and C- terminal configurations), the FBPΔLUZ module (H3H, HispS, CPH and NPGA) and p19. (b) Confocal microscope characterization of *N. benthamiana* leaf samples infiltrated with DsRed:LUZ or LUZ:DsRed, FBPΔLUZ module, p19 and a battery of subcellular markers targeting Golgi, Endoplasmic reticulum (ER), Plasmatic Membrane (PM), Mitochondria and Chloroplast stroma.

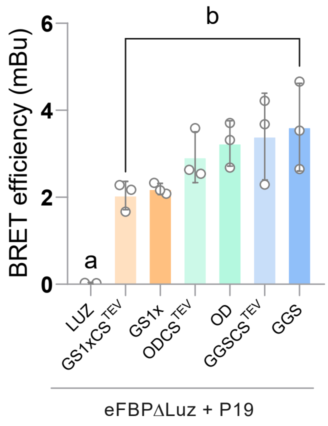


**Supplementary Figure 4.** BRET efficiency of linkers without and with cleavage site calculated with the picture displayed in Figure 3 c.

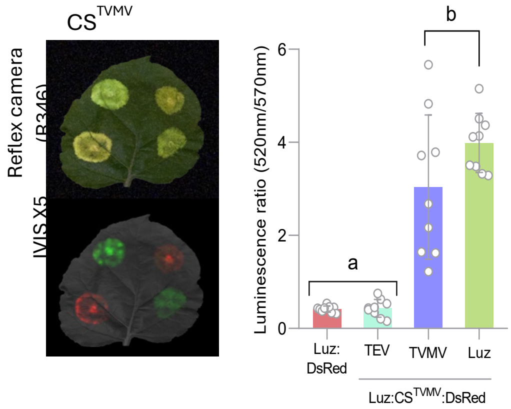


**Supplementary Figure 5.** TVMV-responsive construct displayed equivalent specificity, responding only to TVMV and not to TEV.

Table S1. **List of GoldenBraid plasmids used in this work.** Sequence information is available at https://goldenbraidpro.com/search/features/ by entering the GB number.

| GB number | Construct | Description | *A. tumafaciens* strain |
|---|---|---|---|
| GB5058 | BeYDV:ASCL | Transcriptional unit of Bean Yellow Dwarf Virus-based replicon expressing ASCL gene (a HispS substitute) | EHA105 |
| GB5059 | BeYDV:HmS | Transcriptional unit of Bean Yellow Dwarf Virus-based replicon expressing HmS gene (a HispS substitute) | EHA105 |
| GB5060 | BeYDV:PKS2 | Transcriptional unit of Bean Yellow Dwarf Virus-based replicon expressing PKS2 gene (a HispS substitute) | EHA105 |
| GB4901 | Rep/RepA – p19 | Module for the expression of the Rep/RepA proteins for BeYDV replication and the p19 silencing suppressor | EHA105 |
| GB5096 | TMV:HmS | Transcriptional unit of Tobacco Mosaic Virus-based replicon expressing HmS gene (a HispS substitute) | EHA105 |
| GB5097 | TMV:ASCL | Transcriptional unit of Tobacco Mosaic Virus-based replicon expressing ASCL gene (a HispS substitute) | EHA105 |
| GB5098 | TMV:PKS2 | Transcriptional unit of Tobacco Mosaic Virus-based replicon expressing PKS2 gene (a HispS substitute) | EHA105 |
| GB5135 | PVX:HmS | Transcriptional unit of Potato Virus X-based replicon expressing HmS gene (a HispS substitute) | EHA105 |
| GB5136 | PVX:ASCL | Transcriptional unit of Potato Virus X -based replicon expressing ASCL gene (a HispS substitute) | EHA105 |
| GB5137 | PVX:PKS2 | Transcriptional unit of Potato Virus X -based replicon expressing PKS2 gene (a HispS substitute) | EHA105 |
| GB5091 | eFBPΔLuz | Module for the expression of N. nambi NPGA, H3H, HISPS and CPH genes | EHA105 |
| GB5482 | 3a1_P35S:LUZ:GS1x:DsRed:TNOS | TU for the expression of LUZ:GS1x:DsRed under the P35S promoter and TNOS. | EHA105 |

| | | | |
|---|---|---|---|
| GB4918 | 3a2_P35S(noATG):DsRed:LUZ:TNOS | TU for the expression of DsRed:GS1X:LUZ under the P35S promoter and TNOS. | EHA105 |
| GB5483 | 3a1_P35S:LUZ:GS3x:DsRed:TNOS | TU for the expression of LUZ:GS3x:DsRed under the P35S promoter and TNOS. | EHA105 |
| GB5484 | 3a1_P35S:LUZ:OD:DsRed:TNOS | TU for the expression of LUZ:OD:DsRed under the P35S promoter and TNOS. | EHA105 |
| GB5485 | 3a1_P35S:LUZ:GGS:DsRed:TNOS | TU for the expression of LUZ:GGS:DsRed under the P35S promoter and TNOS. | EHA105 |
| GB5486 | 3a1_P35S:LUZ:Acrlinker:DsRed:TNOS | TU for the expression of LUZ:Acrlinker:DsRed under the P35S promoter and TNOS. | EHA105 |
| GB5487 | 3a1_P35S:LUZ:GS1xTEV:DsRed:TNOS | TU for the expression of LUZ:GS1X:TEV:DsRed under the P35S promoter and TNOS. | EHA105 |
| GB5488 | 3a1_P35S:LUZ:GGSTEV:DsRed:TNOS | TU for the expression of LUZ:GGS:TEV:DsRed under the P35S promoter and TNOS. | EHA105 |
| GB5489 | 3a1_P35S:LUZ:ODTEV:DsRed:TNOS | TU for the expression of LUZ:ODTEV:DsRed under the P35S promoter and TNOS. | EHA105 |
| GB5490 | 3a1_P35S:LUZ:GGSTVMV:DsRed:TNOS | TU for the expression of LUZ:GGSTVMV:DsRed under the P35S promoter and TNOS. | EHA105 |
| GB5491 | 3a2_P35S:LUZ:PdaC1:TNOS | TU for the expression of LUZ:PdaC1 under the P35S promoter and TNOS | EHA105 |
| GB5502 | 3a1_P35S:PdaC1:LUZ:TNOS | TU for the expression of LUZ fused PdaC1 as Ntag | EHA105 |
| GB0594 | TEV NIaPro | TU for the constitutive expression of the Tobacco Etch Virus NIa protease under the P35S promoter and TNOS | GV3101 |
| GB5748 | TVMV NiaPro | TU for the constitutive expression of the Tobacco Vein Mottling Virus NIa protease under the P35S promoter and TNOS | GV3101 |

| | | | |
|---|---|---|---|
| GB1203 | P19 | TU for the expression of the silencing suppressor P19 under the P35S promoter and TNOS | GV3101 |
| GB1236 | EV | Empty vector ( Twister plasmid to swap inserts from an alpha1 or alpha1R vector to any omega level vector) | GV3101 |
| GB3477 | Luz | TU of Luz gene from N. nambi codon optimized for N.benthamiana under the P35S promoter and TNOS | GV3101 |
| GB5510 | HygR- LUZ:GGS-TEV:DsRed | Module for the expression of plXB2alpha1:HygR- Disct02-LUZ:GS1xTEV:DsRed-Disct02. | LBA4404 |
| GB5511 | HygR- LUZ:ODTEV:DsRed | Module for the expression of plXB2alpha1:HygR-Disct02-LUZ:ODTEV:DsRed- Disct02 | LBA4404 |
| GB5108 | nptII-FBPΔLuz | Module for the expression of nptII and the N. nambi NPGA, H3H, HISPS and CPH genes flanked by insulators | LBA4404 |
| GB5512 | HygR-Luz-DsRed (GGS-TVMV) | Module for the expression of plXB2alpha1:HygR- Disct02-LUZ:GGSTVMV:DsRed-Disct02 in PLX vector. | LBA4404 |

**Table S2**. Primers used in this work

| Primer ID | Sequence | Purpose |
|---|---|---|
| MR25JUN01_PVX_FW | ttcaggcctgttcaccatcc | Recombinant PVX systemic detection |
| CC21MAR29_PVX_RV | tggtggtggtagagtgacaac | Recombinant PVX systemic detection |
| MR25JUN02_TMV_FW | aagatgaagccgagacatcgg | Recombinant TMV systemic detection |
| MR25JUN03_TMV_RV | gctccaagacactaccctttcg | Recombinant TMV systemic detection |
| D2336_PVX_FW | atgtcagcaccagctagcac | Wild-type PVX systemic detection |
| D2410_PVX_RV | tggtggtggtagagtgacaac | Wild-type PVX systemic detection |
| D3778_TMV_FW | tgtcttacagtatcactactcca | Wild-type TMV systemic detection |
| D4382_TMV_RV | cctaacagtgctgtgactagc | Wild-type PVX systemic detection |
| D3599_TEV_FW | catctgtgcatcaatgatcgaa | TEV systemic detection |
| D3600_TEV_RV | gtgtggctcgagcatttgacaa | TEV systemic detection |
| MV25APR01_TVMV_FW | tgccatatcaccagaacttgaac | TVM systemic detection |
| MV25APR02_TVMV_RV | cttcccagcatctactgtatcac | TVM systemic detection |

**Table S3.** List of Sub-cellular marker plasmids used in this work.

| Codename | Name | *Agrobacterium* strain used |
|---|---|---|
| Cis-Golgi | pCBNoxbaIGman-GFP cisgolgi | GV3101 |
| Trans-Golgi | pCBNoxbaI STtmd-YFP transgolgi | GV3101 |
| ER | pMOG ER GFP | GV3101 |
| Plasmatic Membrane (PM) | pCB NOX-GFP rapGatPasa (membrana plasmática) | GV3101 |
| Mitochondria (M) | pBINmGFP ATPasa (mitocondria) | GV3101 |
| Stroma (St) | pMOG AtSALI GFP (cloroplasto estroma) | GV3101 |

### Methods S1

The tables provided below show the Agrobacterium cultures co-infiltrated in the same mix for each experiment.

*Assays for LUZ-DsRed BRET pair characterization (Figure 1a-f)*

| ***Construct*** | **EV** | **Luz** | **DsRed:Luz** | **Luz:DsRed** |
|---|---|---|---|---|
| GB1203 | + | - | - | - |
| GB1236 | + | + | + | + |
| GB5091 | + | + | + | + |
| GB3477 | - | + | - | - |
| GB4918 | - | - | + | - |
| GB5482 | - | - | - | + |

*Assays for LUZ-PdaC1 BRET pair characterization (Figure 1e-f)*

| ***Construct*** | **EV** | **Luz** | **PdaC1:Luz** | **Luz:PdaC1** |
|---|---|---|---|---|
| GB1203 | + | - | - | - |
| GB1236 | + | + | + | + |
| GB5091 | + | + | + | + |
| GB3477 | - | + | - | - |
| GB5502 | - | - | + | - |
| GB5491 | - | - | - | + |

*Assays for LUZ-DsRed BRET linker characterization (Figure 1g-i)*

| ***Construct*** | **Luz** | **GS1x** | **GS3x** | **OD** | **GGS** | **AcrLinker** |
|---|---|---|---|---|---|---|
| GB1203 | + | + | + | + | + | + |
| GB5091 | + | + | + | + | + | + |
| GB3477 | + | | - | - | - | - |
| GB5482 | - | + | - | - | - | - |
| GB5483 | - | - | + | - | - | - |
| GB5484 | - | - | - | + | - | - |
| GB5485 | - | - | - | - | + | - |
| GB5486 | - | - | - | - | - | + |

*Assay for LUZ-DsRed BRET pairs with TEV cleavage site BRET efficiency characterization (Figure 2b)*

| ***Construct*** | **Luz** | **GS1x** | **GS1xCS TEV** | **OD** | **ODCS TEV** | **GGS** | **GGSCS TEV** |
|---|---|---|---|---|---|---|---|
| GB1203 | + | + | + | + | + | + | + |
| GB5091 | + | + | + | + | + | + | + |
| GB3477 | + | - | - | - | - | - | - |
| GB5482 | - | + | - | - | - | - | - |
| GB5487 | - | - | + | - | - | - | - |
| GB5484 | - | - | - | + | - | - | - |
| GB5489 | - | - | - | - | + | - | - |
| GB5486 | - | - | - | - | - | + | - |
| GB5488 | - | - | - | - | - | - | + |

*Assay for LUZ-DsRed BRET pairs with TEV cleavage for TEV protease recognition (Figure 2c-e)*

| | | **GS1x** | | **OD** | | **GGS** | |
|---|---|---|---|---|---|---|---|
| ***Construct*** | **Luz** | **-TEVPro** | **+TEVPro** | **-TEVPro** | **+TEVPro** | **-TEVPro** | **+TEVPro** |
| GB1203 | + | + | + | + | + | + | + |
| GB5091 | + | + | + | + | + | + | + |
| GB3477 | + | - | - | - | - | - | - |
| GB0594 | - | - | + | - | + | - | + |
| GB5487 | - | + | + | - | - | - | - |
| GB5489 | - | - | - | + | + | - | - |
| GB5488 | - | - | - | - | - | + | + |

*Assay for LUZ-DsRed BRET pair with TEV cleavage for subcellular localization under TEV protease treatment (Figure 2f)*

| | **Cis-Golgi** | | **Trans-Golgi** | | **ER** | | **Plasmatic Membrane** | | **Mitochondria** | | **Stroma** | |
|---|---|---|---|---|---|---|---|---|---|---|---|---|
| ***Construct*** | **-TEVPro** | **+TEVPro** | **-TEVPro** | **+TEVPro** | **-TEVPro** | **+TEVPro** | **-TEVPro** | **+TEVPro** | **-TEVPro** | **+TEVPro** | **-TEVPro** | **+TEVPro** |
| GB1203 | + | + | + | + | + | + | + | + | + | + | + | + |
| GB5091 | + | + | + | + | + | + | + | + | + | + | + | + |
| GB0594 | - | + | - | + | - | + | - | + | - | + | - | + |
| GB5488 | + | + | + | + | + | + | + | + | + | + | + | + |
| Cis-Golgi | + | + | - | - | - | - | - | - | - | - | - | - |
| Trans-Golgi | - | - | + | + | - | - | - | - | - | - | - | - |
| ER | - | - | - | - | + | + | - | - | - | - | - | - |
| Plasmatic Membrane | - | - | - | - | - | - | + | + | - | - | - | - |
| Mitochondria | - | - | - | - | - | - | - | - | + | + | - | - |
| Stroma | - | - | - | - | - | - | - | - | - | - | + | + |

*Assays (Figure 2h-i)*

| | | | Luz:CSTEV:DsRed | | | Luz:CSTVMV:DsRed | | |
|---|---|---|---|---|---|---|---|---|
| ***Construct*** | **Luz** | **Luz:Ds Red** | **EV** | **TVMV** | **TEV** | **EV** | **TEV** | **TVMMV** |
| GB1203 | + | + | + | + | + | + | + | + |
| GB1236 | - | - | + | - | - | + | - | - |
| GB5091 | + | + | + | + | + | + | + | + |
| GB3477 | + | - | - | - | - | - | - | - |
| GB5485 | - | + | + | + | - | - | - | - |
| GB5488 | - | - | + | + | + | - | - | - |
| GB5490 | - | - | - | - | - | + | + | + |
| TEV | - | - | - | - | + | - | + | - |
| TVMV | - | - | - | + | - | - | - | + |

*Assay for LUZ-DsRed and Luz-PdaC1 BRET pairs characterization with a reflex camera system coupled with filters (Figure 2)*

| ***Construct*** | **EV** | **Luz** | **DsRed:Luz** | **Luz:DsRed** | **PdaC1:Luz** | **Luz:PdaC1** |
|---|---|---|---|---|---|---|
| GB1203 | + | - | - | - | - | - |
| GB1236 | + | + | + | + | + | + |
| GB5091 | + | + | + | + | + | + |
| GB3477 | - | + | - | - | - | - |
| GB4918 | - | - | + | - | - | - |
| GB5482 | - | - | - | + | - | - |
| GB5502 | - | - | - | - | + | - |
| GB5491 | - | - | - | - | - | + |

*Assay for LUZ-DsRed BRET pair subcellular localization (Figure S2)*

| | Cis-Golgi | | Trans-Golgi | | ER | | Plasmatic Membrane | | Mitochondria | | Stroma | |
|---|---|---|---|---|---|---|---|---|---|---|---|---|
| ***Construct*** | **LUZ** | **+TEVPro** | **-TEVPro** | **+TEVPro** | **-TEVPro** | **+TEVPro** | **-TEVPro** | **+TEVPro** | **-TEVPro** | **+TEVPro** | **-TEVPro** | **+TEVPro** |
| GB1203 | + | + | + | + | + | + | + | + | + | + | + | + |
| GB5091 | + | + | + | + | + | + | + | + | + | + | + | + |
| GB3477 | + | - | + | - | + | - | + | - | + | - | + | - |
| GB0594 | - | + | - | + | - | + | - | + | - | + | - | + |
| Cis-Golgi | + | + | - | - | - | - | - | - | - | - | - | - |
| Trans-Golgi | - | - | + | + | - | - | - | - | - | - | - | - |
| ER | - | - | - | - | + | + | - | - | - | - | - | - |
| Plasmatic Membrane | - | - | - | - | - | - | + | + | - | - | - | - |
| Mitochondria | - | - | - | - | - | - | - | - | + | + | - | - |
| Stroma | - | - | - | - | - | - | - | - | - | - | + | + |
